# Thiol efflux mediated by an ABCC-like transporter participates for *Trypanosoma cruzi* adaptation to environmental and chemotherapeutic stresses

**DOI:** 10.1101/2020.03.26.009753

**Authors:** Kelli Monteiro da Costa, Eduardo J. Salustiano, Raphael do Carmo Valente, Leonardo Freire-de-Lima, Lucia Mendonça-Previato, José Osvaldo Previato

**Affiliations:** Laboratório de Glicobiologia, Instituto de Biofísica Carlos Chagas Filho, Universidade Federal do Rio de Janeiro - UFRJ, Rio de Janeiro/RJ, Brasil; Faculdade de Ciências Médicas, Universidade do Estado do Rio de Janeiro, Rio de Janeiro, Brasil

**Keywords:** ABC transporter, benznidazole, ceramide, drug resistance, iron, MRP-1, P-glycoprotein, thiol, *Trypanosoma cruzi*, oxidative stress

## Abstract

The protozoan *Trypanosoma cruzi* is the etiologic agent for Chagas disease, which affects 6-7 million people worldwide. The parasite presents high biological diversity, reflecting on the inefficiency of benznidazole in chronic or older patients. ABC superfamily proteins contain active transporters involved in the xenobiotic and endobiotic efflux and overexpressed in MDR cells. An ABCC-like transport was identified in the *T. cruzi* Y strain, being able to extrude thiol-conjugated compounds. As non-protein thiols represent prime line of defense towards reactive species, ABCC-like activity could participate in the regulation of mediators implicated in responses to cellular stress arising from a variety of stimuli, as environmental or chemotherapeutic. This study shows that *T. cruzi* ABCC-like protein transports GSH, GSSG and ceramides, all implicated in cellular stress. Hemin, representative from the hematophagous feeding of the vector, was transported as well, suggesting a role for ABCC as a metal-thiol transporter. In addition, all strains evaluated showed ABCC-like activity, while no ABCB1-like activity was detected. Also, results suggest that ABCC-like does not associate to natural resistance to benznidazole, considering that the sensitive strains CL Brener and Berenice showed higher ABCC-like activity than the resistant strains Y and Colombiana. Instead, ABCC-like efflux increased after continuous exposure of Y strain to benznidazole. Moreover, ABCC does not perform direct efflux of drug and its participation in the machinery of protection against stress depends on the efflux of metabolites in conjugation to or in cotransport with thiol.

## Introduction

*Trypanosoma cruzi* is a flagellated protozoan responsible by causing anthropozoonosis Chagas disease (1). According to the World Health Organization, Chagas disease is classified as a neglected tropical disease with an estimative of 6-7 million infected people in the world and almost 65 million at risk of infection. The Pan American Health Organization defined as endemic area from southern United States to southern Argentina and Chile, totaling 21 countries (2). The main form of transmission is through the insect vector, predominant in rural areas with rudimentary infrastructure, promoting a favorable environment for its reproduction as well as proximity to the wild cycle (3). Alternatively, *T. cruzi* may be transmitted by the consumption of food contaminated with vector feces or with the secretion of infected mammals (4), donation of blood, organs or tissues, especially in countries that do not screen samples for *T. cruzi* (5), and during pregnancy (6). Furthermore, sexual transmission (7, 8) and other species as potential vectors (9) show up as possible routes for spreading of disease mainly in non-endemic areas.

First-line treatment of Chagas disease is performed with benznidazole, whose success depends on the stage of the disease, the patient’s age, and on biochemical characteristics of the strain (10). *T. cruzi* strains show great discrepancy in susceptibility to chemotherapy, being resistance detected on wild-type strains or after prolonged treatment (11–13). Resistance to benznidazole in *T. cruzi* is multifactorial and results from mechanisms involving prodrug activation, defenses against free radicals and drug efflux (14).

The life cycle of *T. cruzi* is complex and involves the epimastigote and amastigote replicative stages in the invertebrate and vertebrate hosts respectively, and the trypomastigote infective stage (15). In the hematophagous invertebrate host, parasites must thrive in a pro-oxidant microenvironment caused by the blood degradation and heme release (16). Regardless of being necessary as cofactor for several biological processes in the parasite (17), heme has cytotoxic action due to the generation of reactive oxygen species (ROS) (18). Even though most ROS are a consequence of cellular respiration, xenobiotics present an important source of oxidative stress, either releasing ROS by biotransformation or via direct consumption of antioxidant defenses (19). In addition, stress-inducing agents are able to affect sphingolipid metabolism, resulting in the accumulation of ceramides (20), which is a sphingolipid comprised of a sphingosine-related base linked to a fatty acid through an amide bond. Ceramides are ubiquitous in nature as components of cell membranes and as regulators of cell cycle arrest, differentiation, cell senescence and apoptosis (21). The nature of the ceramide-mediated responses suggests that this sphingolipid coordinates response pathways to intracellular stresses (22), since its production is sensitive to ROS levels (23).

ABCC is an ABC active transporter subfamily, studied in several organisms owing to their ability to extrude endobiotics or xenobiotics alone or in conjugation to or in cotransport with phosphate, glucuronide or GSH (24). Human ABCC1 is able to transport sphingolipids as sphingosine-1-phosphate (25), sphingomyelin and glucosylceramides (26). Moreover, ABCC members are involved with chemotherapy resistance phenotype in major protozoan parasites including *Leishmania*, *Trypanosoma* and *Plasmodium* species (27). In *T. cruzi*, 27 ABC genes were identified in the genome, the first being named PGP1 and PGP2 by Dallagiovanna *et al.* (28, 29). Subsequently, it was observed that the PGP1 and PGP2 genes show great homology with the ABCC6 and ABCC2 genes of *L. major* and *T. brucei*, respectively (30). Additionally, *T. cruzi* Y strain showed efflux of thiol-conjugated compounds (31) in a similar mechanism to that performed by ABCC members present on humans (24) as well as on other trypanosomatids (27). Considering the importance of oxidative stress to *T. cruzi* development and the crucial participation of ABC transporters on cellular detoxification, we investigated the efflux of metabolites involved in oxidative stress pathways elicited by the microenvironment and by chemotherapy as well as the importance of ABCC-like activity for *T. cruzi* strains with diverse degrees of resistance to benznidazole.

## Results

### An ABCC-like transporter mediates efflux of GSH and GSSG

Previous results have shown that ABCC was capable of transporting thiol-conjugated compounds in epimastigote forms of *T. cruzi* Y strain (31). Similarly, GSH, GSSG and hemin increased the accumulation of carboxyfluorescein (CF), a fluorescent substrate transported by ABCC subfamily members (Fig. S1). In Fig. 1A and 1B, either reduced (GSH) or oxidized (GSSG) glutathione was able to increase the median fluorescence intensities (MFI) for CF and the percentage of CF-positive parasites, demonstrating that both molecules competitively inhibited CF efflux in the active transport mediated by ABCC. Analyzing the transport inhibition index (Δ), calculated as the ratio of CF MFI in the presence of GSH or GSSG to the one in its absence (control/CTL), it was observed that the reduced peptide inhibited the CF efflux more effectively than its oxidized version (Fig. 1C), suggesting that ABCC has a greater affinity for GSH than GSSG.

**Figure 1.**
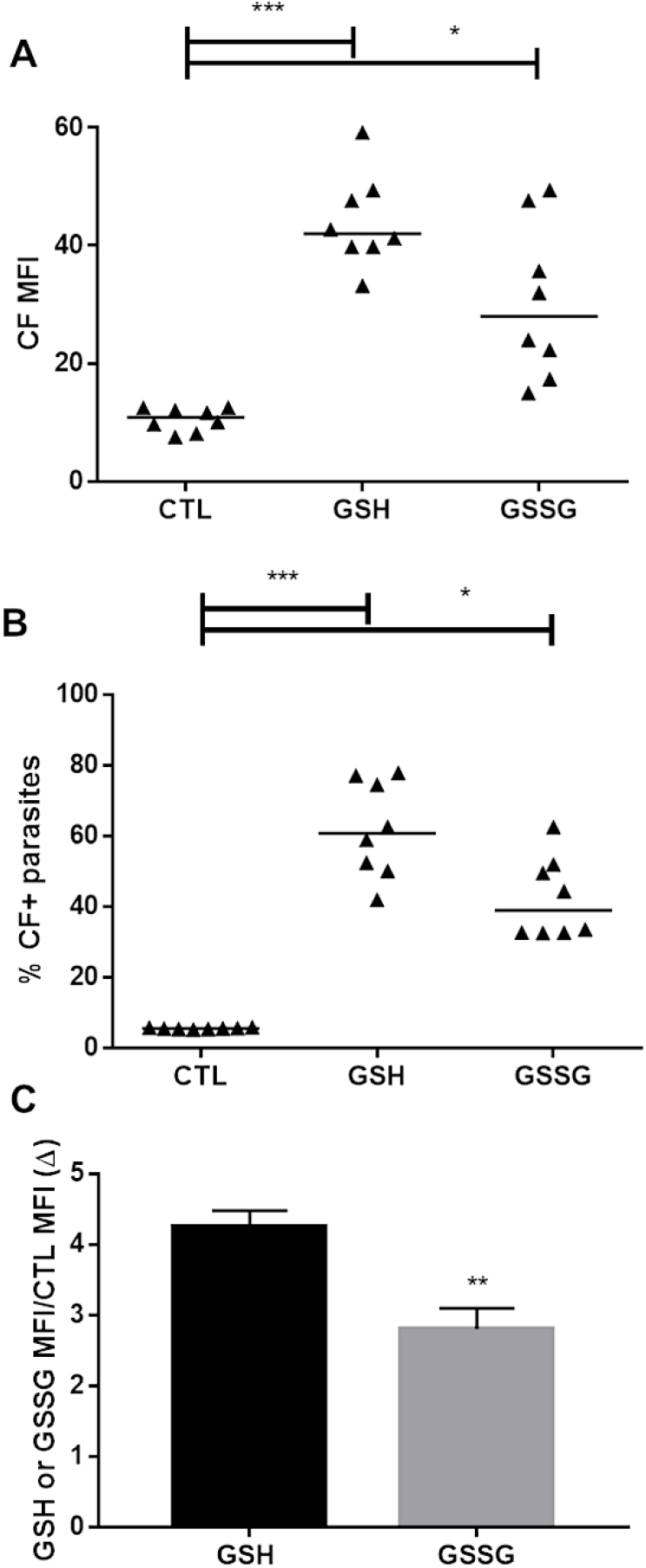
GSH and GSSG modulate CF efflux. ABCC-like activity was evaluated by the carboxyfluorescein (CF) efflux assay. 10^7^ epimastigotes/mL were incubated in medium containing 50 μM 5(6)-carboxyfluorescein diacetate (CFDA) in absence (control, CTL) or presence of 5 mM GSH or GSSG for 30 min. Parasites were then centrifuged and incubated in fresh medium in absence or presence of the modulators for another 30 min. At the end, parasites were centrifuged, suspended in PBS and kept on ice until acquisition by flow cytometry. (A) CF median fluorescence intensities (MFI), (B) percentage of CF+ parasites and (C) Δ values at 27 °C for epimastigote forms of Y strain. Lines represent the median, bars represent mean + SEM and the values of significance were represented by (*) for p<0.05, (**) p <0.01 and (***) p<0.001, n=8 independent experiments.

### Hemin preincubation promotes thiol depletion and CF accumulation

Hematophagous feeding of the invertebrate host results in the release of iron ions in the microenvironment where the epimastigote forms of *T. cruzi* are found. To assess whether the hemin breakdown is able to reduce the parasite’s antioxidant defenses, GSH levels were measured using the fluorescent probe 5-chloromethylfluorescein (CMFDA), which covalently binds with thiol to form the thiol-conjugated methylfluorescein (TMF). Trypanothione [T(SH)_2_] is the main non-protein thiol of the *T. cruzi*, containing two GSH molecules linked by a spermidine chain (32). Thus, the level of TMF would be an indirect measure of GSH and, consequently, of T(SH)_2_. As expected, preincubation with 200 μM hemin, a protoporphyrin containing a ferric iron with a chlorine ligand, reduced TMF levels by 82.50% in epimastigote forms of Y strain (Fig. 2A). The alkylating agent N-ethylmaleimide (NEM), positive control to thiol depletion, reduced TMF levels by 91% compared to control.

**Figure 2.**
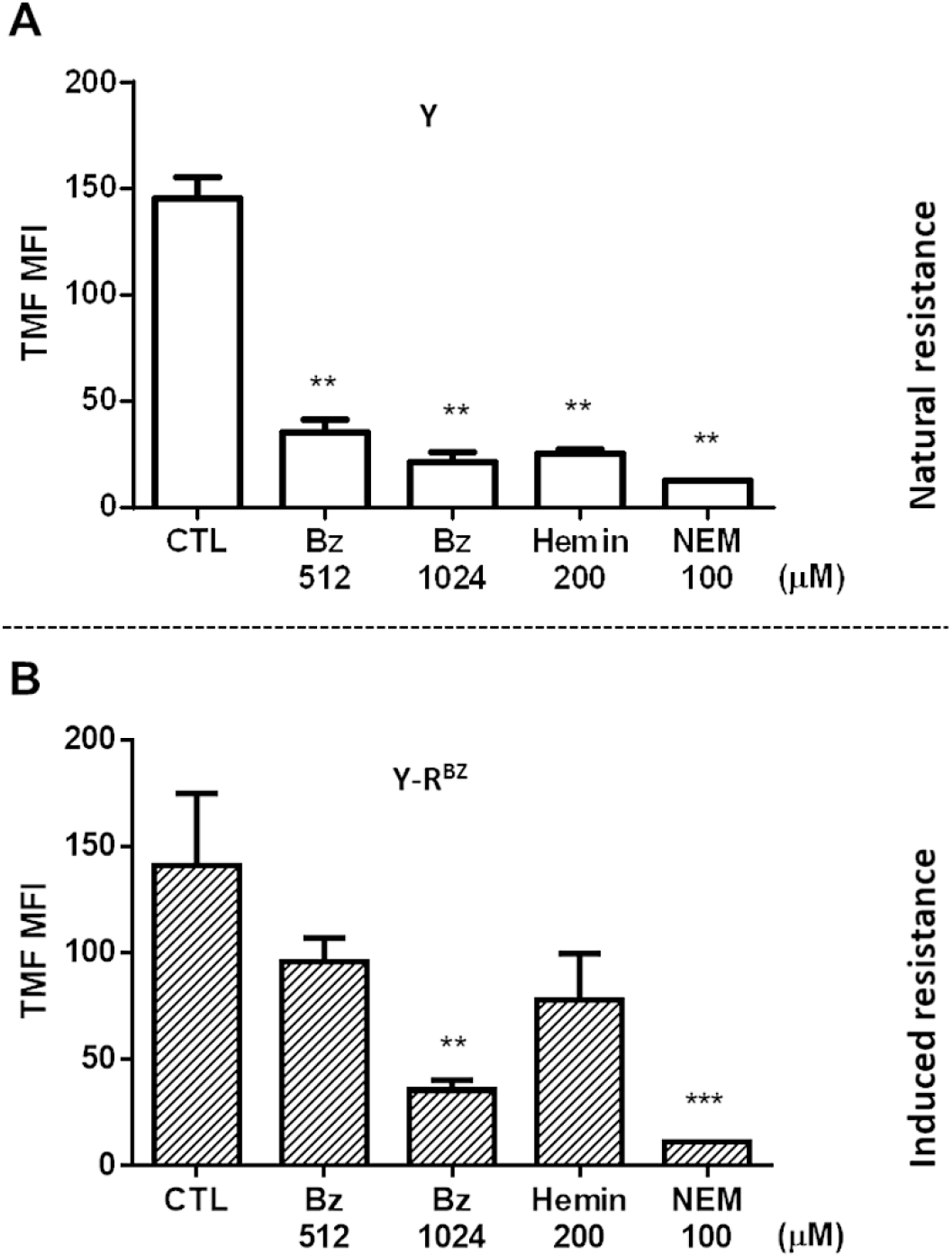
Benznidazole and hemin reduce intracellular levels of free thiols. 10^7^ epimastigotes/mL were incubated in PBS in absence (control, CTL) or presence of 512 or 1024 μM benznidazole or 200 μM hemin for 3 h at 27 °C. Parasites were centrifuged and then incubated in PBS with 1.5 μM 5-chloromethylfluorescein (CMFDA) for 15 min. Next, parasites were kept on ice until acquisition by flow cytometry. 100 μM N-ethylmaleimide (NEM) was used as positive control for thiol depletion. Bars represent the mean + SEM of median fluorescence intensities (MFI) of thiol-conjugated methylfluorescein (TMF) in epimastigote forms of (A) Y strain and (B) Y-R^BZ^ parasites. The values of significance were represented by (**) for p<0.01 and (***) p<0.001, n=3 (Y) and n=5 (Y-R^BZ^) independent experiments.

Preincubation with 200 μM hemin before the ABCC-mediated efflux assay also enabled an increase in CF MFI and in the percentage of CF-positive parasites when the assay was performed at 27 °C (Fig. 3A and 3C) and 37 °C (Fig. S2A and S2C). To verify if hemin could be a direct substrate for ABCC, it was employed as a competitive inhibitor during the CF efflux (Fig. 3B and 3D; Fig. S2B and S2D). In the latter scenario, hemin appears to be less effective in promoting CF accumulation when compared to its effects after preincubation both at 27 °C and 37 °C (Fig. 3E and 3F, respectively). This trend becomes clear when we analyze the Δ values in the assay performed at 37 °C (Fig. 3F). In this case, the Δ value of hemin was higher in the preincubation than in the competitive inhibition. The representative histograms of CF fluorescence when hemin was employed are shown in Fig. S1.

**Figure 3.**
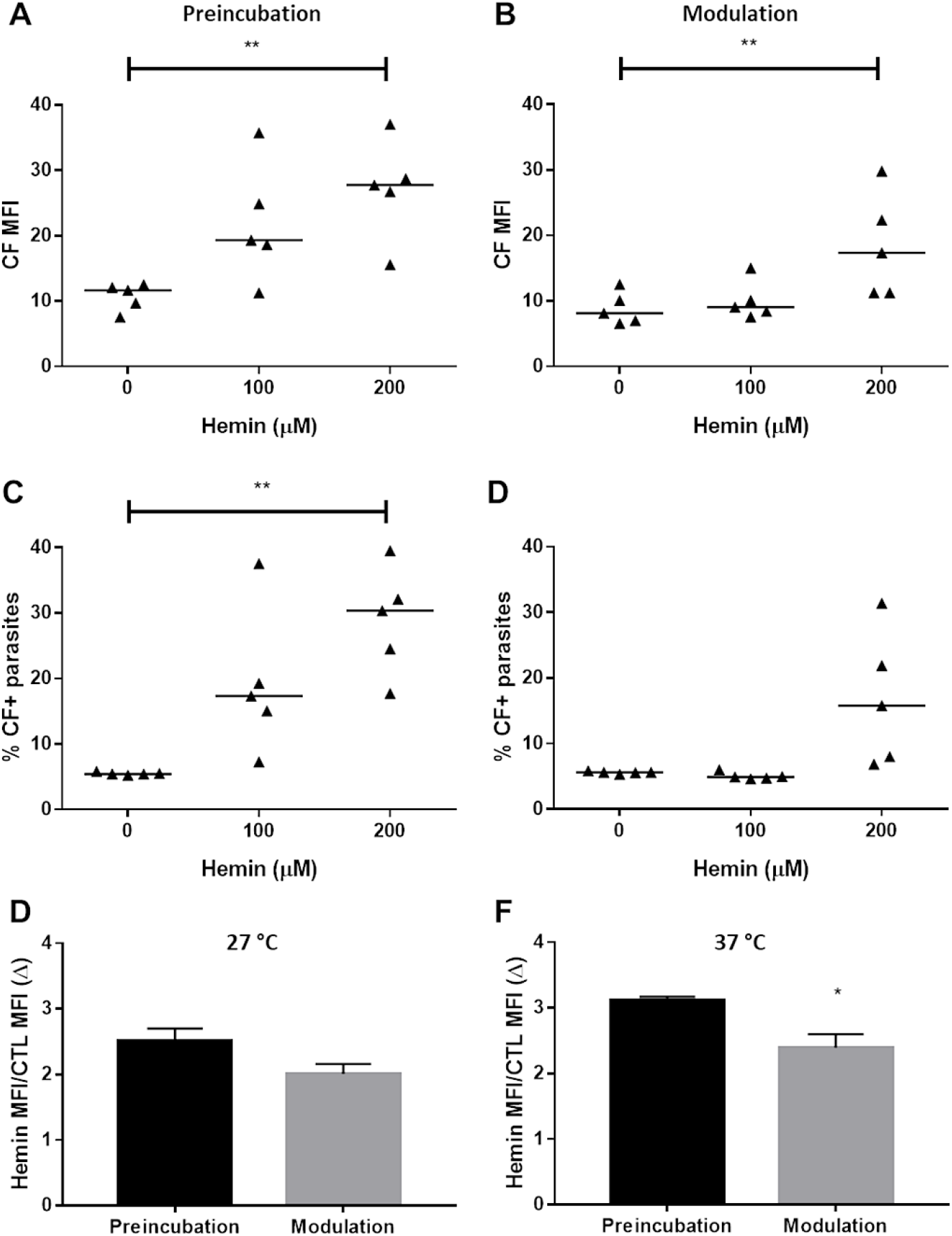
Hemin increases CF accumulation. ABCC-like activity was evaluated by the carboxyfluorescein (CF) efflux assay. 10^7^ epimastigotes/mL were incubated in medium containing 50 μM 5(6)-carboxyfluorescein diacetate (CFDA) in absence (control, CTL) or presence of 100 or 200 μM hemin for 30 min. Parasites were then centrifuged and incubated in fresh medium in absence or presence of hemin for another 30 min. Alternatively, a preincubation in absence (control, CTL) or presence of hemin was performed for 1 h, but in this case no hemin was added in CF efflux assay (A and C). In this case, no hemin was added in efflux assay. At the end, parasites were centrifuged, suspended in PBS and kept on ice until acquisition by flow cytometry. (A and B) CF median fluorescence intensities (MFI), (C and D) percentages of CF+ parasites and Δ values for 200 μM hemin in the assay performed (E) at 27 °C and (F) 37 °C in epimastigote forms of Y strain. (A to D) CF efflux assay at 27 °C, in which lines represent the median. (E and F) Bars represent mean + SEM and the values of significance represented by (*) for p<0.05 and (**) p <0.01, n=5 independent experiments.

### ABCC-like participates of ceramide efflux

Several approaches suggest an important role for sphingolipids in regulating stress responses including apoptosis (23, 33). Knowing that stress stimuli can result in biosynthesis or remodeling of ceramides (34) and that ABC proteins are involved with the transport of sphingolipids (35, 36), we evaluated whether ABCC transporters could mediate the transport of ceramides in *T. cruzi*. For this purpose, a preincubation with sphingosine, which is metabolized by ceramide synthase promoting the accumulation of ceramides (37), was performed prior to the CF efflux assay. In epimastigote forms of Y strain, preincubation with sphingosine increased CF MFI from 10.49 to 63.55 (Fig. 4A). In addition, about 55% of the parasites have accumulated CF in the cytosol (Fig. 4B).

**Figure 4.**
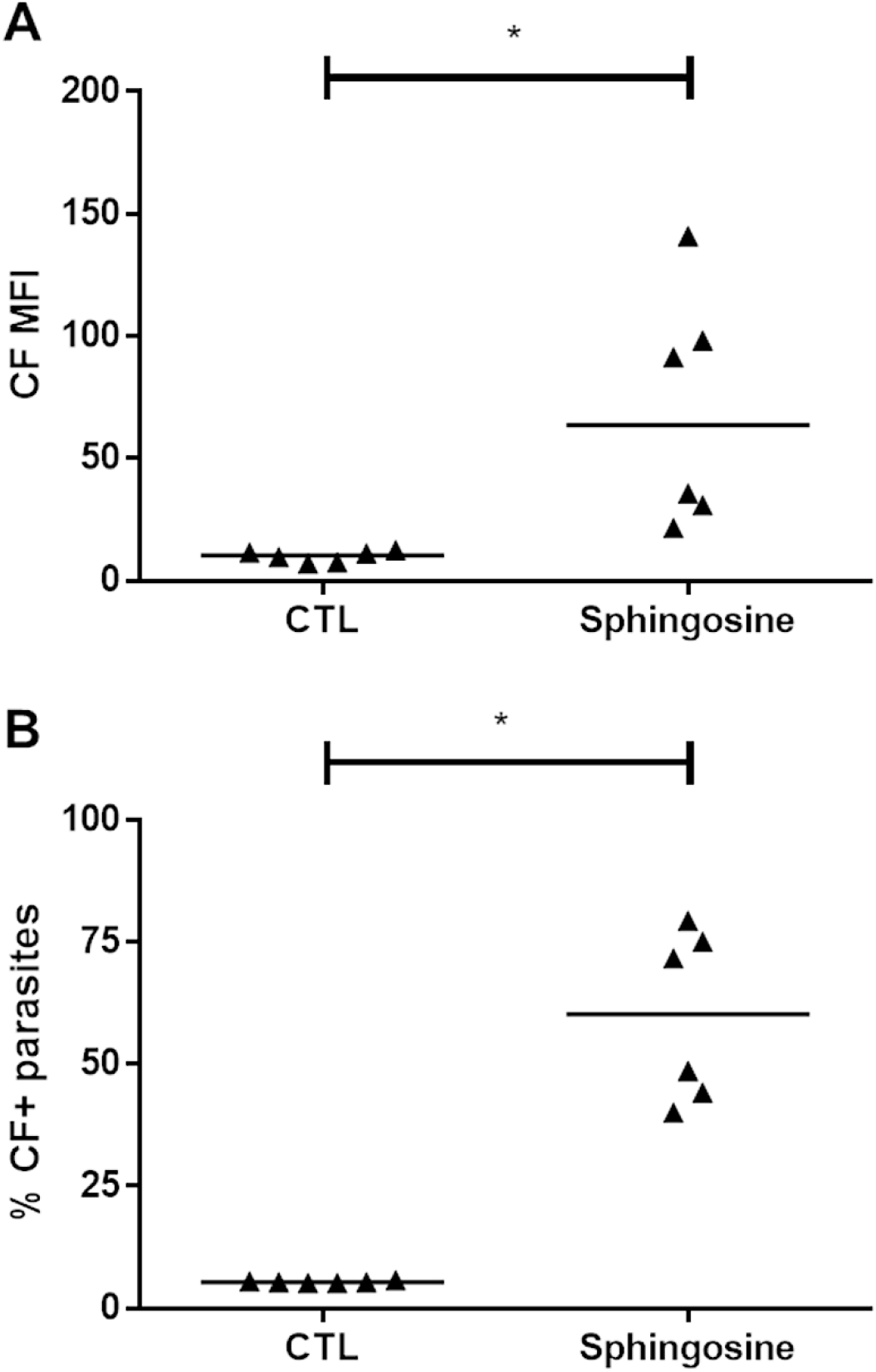
Sphingosine preincubation increases CF accumulation. ABCC-like activity was evaluated by the carboxyfluorescein (CF) efflux assay. 10^7^ epimastigotes/mL were preincubated in medium in absence (control, CTL) or presence of 60 μM sphingosine for 1 h. Parasites were then centrifuged and incubated in fresh medium containing 50 μM 5(6)-carboxyfluorescein diacetate (CFDA) for 30 min. Again, parasites were centrifuged and incubated in fresh medium for another 30 min. At the end, parasites were centrifuged, suspended in PBS and kept on ice until acquisition by flow cytometry. (A) CF median fluorescence intensities (MFI) and (B) percentages of CF+ parasites at 27 °C in epimastigote forms of Y strain. Lines represent the median and values of significance were represented by (*) for p<0.05, n=6 independent experiments.

To rule out the possibility that the results obtained would be by direct transport of sphingosine, the ABCC-mediated efflux assay was carried out with N-[6-[(7-nitro-2-1,3-benzoxadiazol-4-yl)amino]hexanoyl]-D-erythro-sphingosine (C6-NBD-cer), a fluorescent analogue of a biologically active, short-chain membrane-permeable ceramide as substrate. Fig. 5A and 5B show that MK-571, a specific ABCC inhibitor, promoted an increase in C6-NBD-cer MFI with almost 100% of parasites inhibited. ABC proteins rely on ATP to efflux several chemically unrelated molecules and ABCC subfamily members in specific mediate the transport of these molecules in conjugation to or in cotransport with GSH. Therefore, the effect of iodoacetic acid (IAA) and NEM, respectively ATP and thiol depletion agents, were employed in the efflux of fluorescent ceramide. Noteworthy, both agents induced C6-NBD-cer accumulation observed by the increase in MFI and in the percentage of C6-NBD-cer-positive parasites (Fig. 5A and 5B, respectively). However, the percentage of parasites that accumulated the substrate was lower, around 35%. Considering these results, ceramides could be endogenous substrates for ABCC-like transporters in *T. cruzi*. The representative histograms of CF and C6-NBD-cer fluorescence intensities are depicted in Fig. S3.

**Figure 5.**
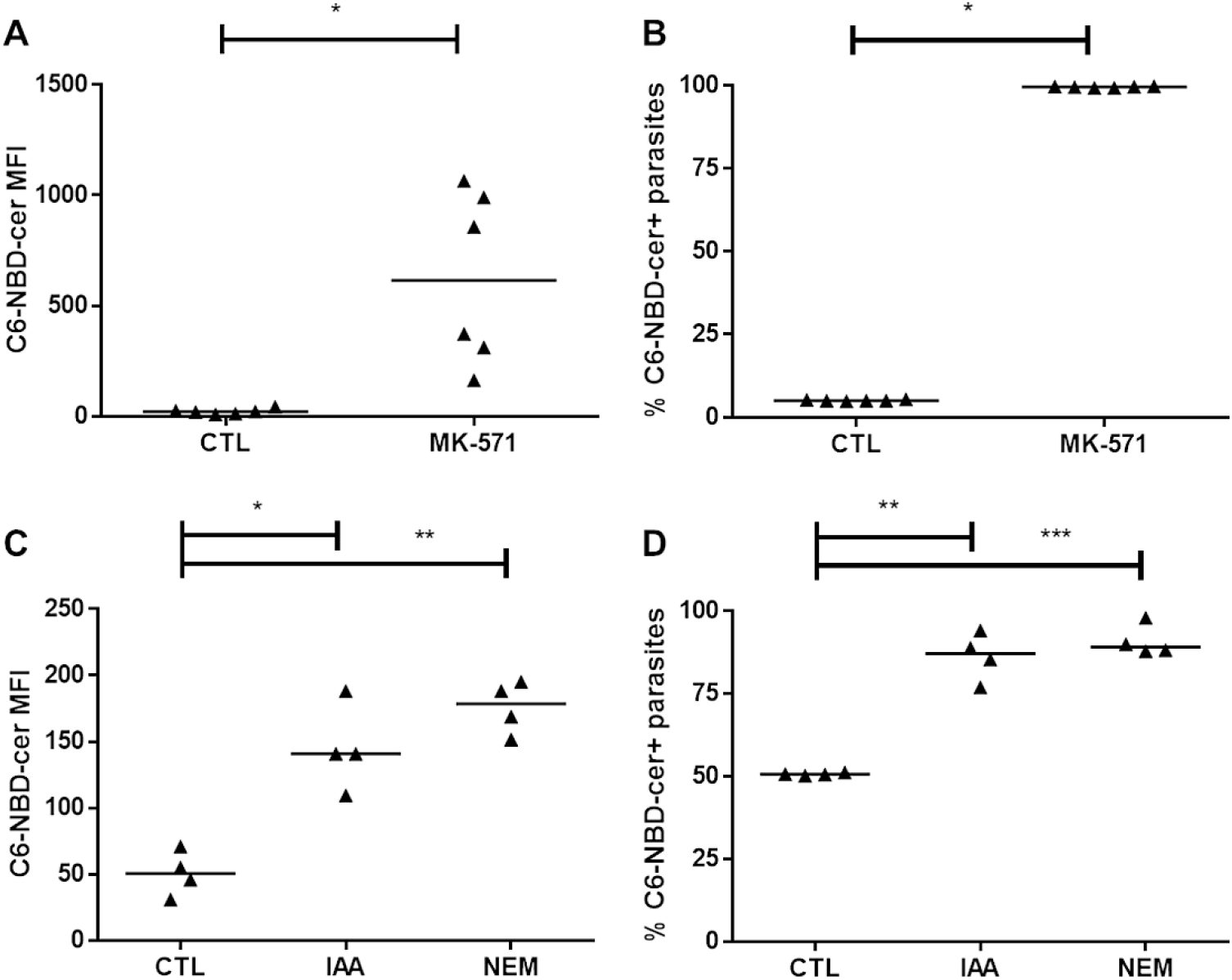
ABCC inhibitor MK-571 and ATP or thiol depletion reduce ceramide efflux. ABCC-like activity was evaluated by the C6-NBD-ceramide (N-[6-[(7-nitro-2-1,3-benzoxadiazol-4-yl)amino]hexanoyl]-D-erythro-sphingosine) efflux assay. 10^7^ epimastigotes/mL were incubated in medium containing 10 μM C6-NBD-cer in absence (control, CTL) or presence of 200 μM MK-571 for 30 min. Parasites were then centrifuged and incubated in fresh medium in absence or presence of the inhibitor for another 30 min. Alternatively, a preincubation in absence (control, CTL) or presence of 2 mM iodoacetic acid (IAA) or 100 μM N-ethylmaleimide (NEM) were employed for ATP or thiol depletion for 1 h and removed before the efflux assay. In this case, neither IAA, NEM or MK-571 was added in the efflux assay. At the end, parasites were centrifuged, suspended in PBS and kept on ice until acquisition by flow cytometry. (A and C) C6-NBD-cer median fluorescence intensities (MFI) and (B and D) percentages of C6-NBD-cer+ parasites at 27 °C for epimastigote forms of Y strain. Lines represent the median and the values of significance were represented by (*) for p<0.05, (**) p <0.01 and (***) p<0.001, n=6 (MK-571) and n=4 (IAA and NEM) independent experiments.

### Resistance to benznidazole

In the present study, four *T. cruzi* strains considered naturally susceptible or resistant to benznidazole were evaluated for CF efflux aptitude. To confirm the susceptibility to benznidazole, epimastigote forms from Berenice, CL Brener, Y and Colombiana strains were treated *in vitro* with a range of concentrations of drug for 48 h. Fig. S4A shows the IC_50_ values to the four strains calculated by mitochondrial reducing activity. Results indicated that Berenice and CL Brener strains are susceptible to treatment, with IC_50_ of 7.06 ± 1.16 μM and 10.42 ± 1.83 μM respectively. The Y and Colombiana strains were significantly more resistant, presenting IC_50_ of 31.02 ± 2.89 μM and 33.90 ± 1.41 μM respectively.

Additionally, epimastigote forms from Y strain were selected *in vitro* after exposure to benznidazole as described in Experimental procedures. This methodology produced great impact on adaptation to chemoterapy, since its IC_50_ reached 395.10 μM (Fig. S4B). From now on, the Y strain selected *in vitro* to benznidazole will be referred to as Y-R^BZ^.

### Absence of ABCB1-like efflux

Our results demonstrated the lack of ABCB1-like activity in epimastigote and trypomastigote forms from Y strain (31). In order to confirm that this transporter is not involved in the results obtained in the present work, the ABCB1-like mediated efflux assay was performed with the fluorescent substrate rhodamine 123 (Rho 123). The Fig. S5 (left panel) displays representative histograms for Rho 123 fluorescence on epimastigote forms from CL Brener, Berenice, Colombiana and Y-R^BZ^. Likewise, neither cyclosporine A (CsA), verapamil (VP) or trifluoperazine (TFP), known ABCB1 inhibitors, increased Rho 123 MFI or the percentage of Rho 123-positive parasites in any of the *T. cruzi* strains (Fig. S6).

### Natural resistance to benznidazole does not relate to ABCC-like mediated efflux

The metabolism of benznidazole is carried out by nitroreductases and proceeds via short-lived intermediates, which could conjugate to free thiols (38). Similar to the effects of hemin, preincubation with benznidazole for 3 h reduced thiol levels by 75.83% and 85.13% at concentrations of 512 μM and 1024 μM respectively when compared to the control (Fig. 2A). As benznidazole reduced thiol levels and ABCC relates to MDR via transport of xenobiotics conjugated to or in cotransport with GSH, the ABCC-like mediated efflux assay was performed in epimastigote forms of naturally resistant (Colombiana) or sensitive strains (Berenice and CL Brener) to the drug. All strains exhibited CF efflux at 27 °C, as shown in the representative histograms for CF fluorescence (Fig. S5 right panel). CF MFI and percentages of CF-positive parasites were higher in the presence of the inhibitors MK-571 (Fig. 6A to 6F) and indomethacin (Fig. S7A to S7F) relative to control. Akin to Y strain, ABCC-mediated transport was higher when the temperature was raised from 27 °C to 37 °C (Fig. S8A to S8F). For comparing ABCC-like mediated efflux among *T. cruzi* strains, the Δ values were calculated for MK-571 since it does not inhibit other ABC subfamilies (Fig. 7A and 7B). At both temperatures, Berenice and CL Brener presented higher Δ values than Y and Colombiana strains, suggesting a greater aptitude for CF efflux in those strains and, consequently, the fact that ABCC inversely correlates to natural resistance to benznidazole. The Δ values of Y strain were obtained from da Costa *et al.* (31) and included for comparison.

**Figure 6.**
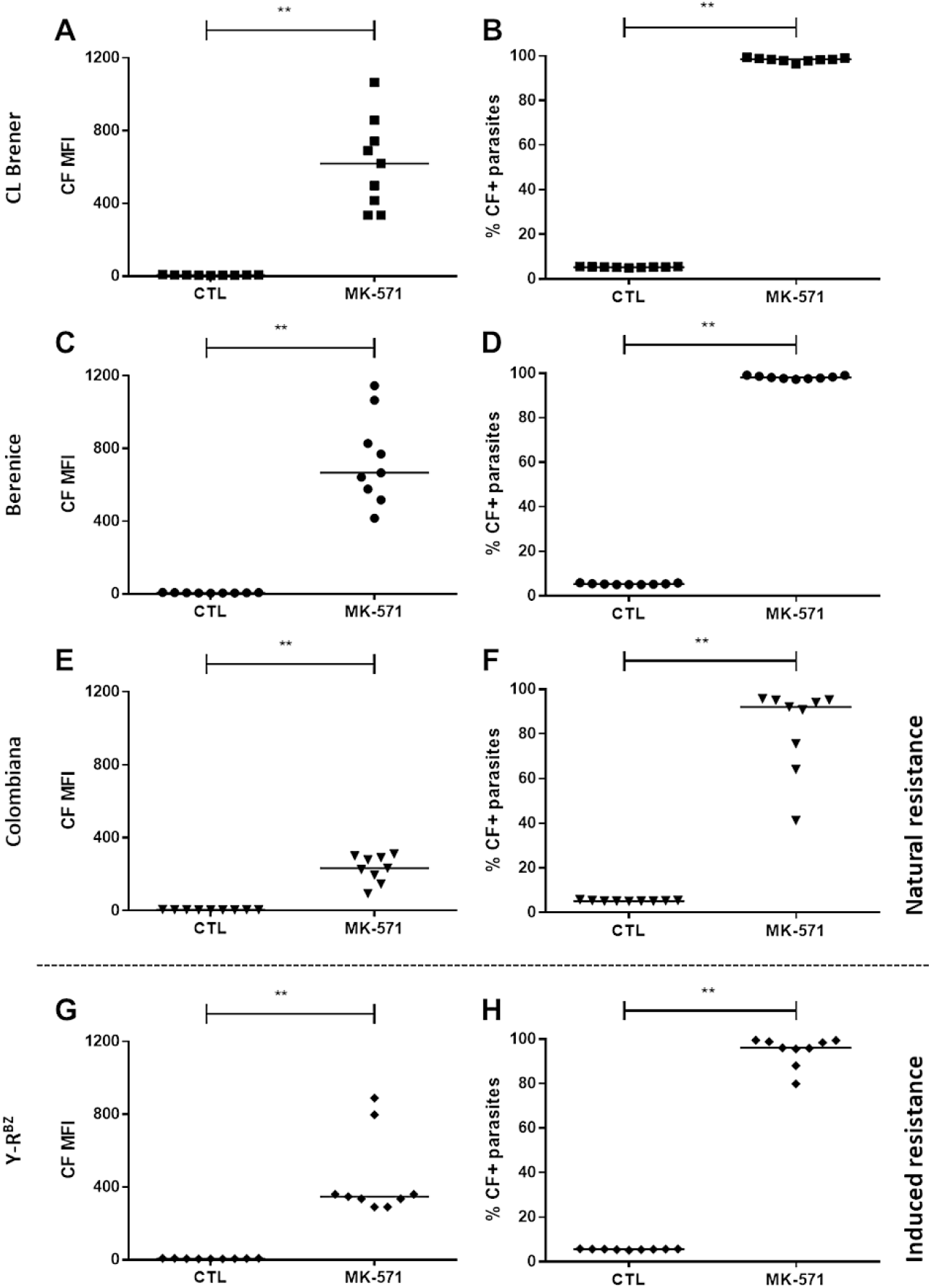
ABCC inhibitor MK-571 increases CF accumulation in *T. cruzi* sensitive and resistant to benznidazole. ABCC-like activity was evaluated by the carboxyfluorescein (CF) efflux assay at 27 °C. 10^7^ epimastigotes/mL were incubated in medium containing 5 μM 5(6)-carboxyfluorescein diacetate (CFDA) in absence (control, CTL) or presence of 200 μM MK-571 for 30 min. Parasites were centrifuged and incubated in medium in absence or presence of the inhibitor for another 30 min. Next, parasites were centrifuged, suspended in PBS and kept on ice until acquisition by flow cytometry. Graphs exhibit CF median fluorescence intensities (MFI) (left panel) and percentages of CF+ parasites (right panel) for epimastigote forms from (A and B) CL Brener, (C and D) Berenice and (E and F) Colombiana strains and (G and H) Y-R^BZ^ parasites. Lines represent the median and values of significance were represented by (**) for p <0.01, n=9 independent experiments.

**Figure 7.**
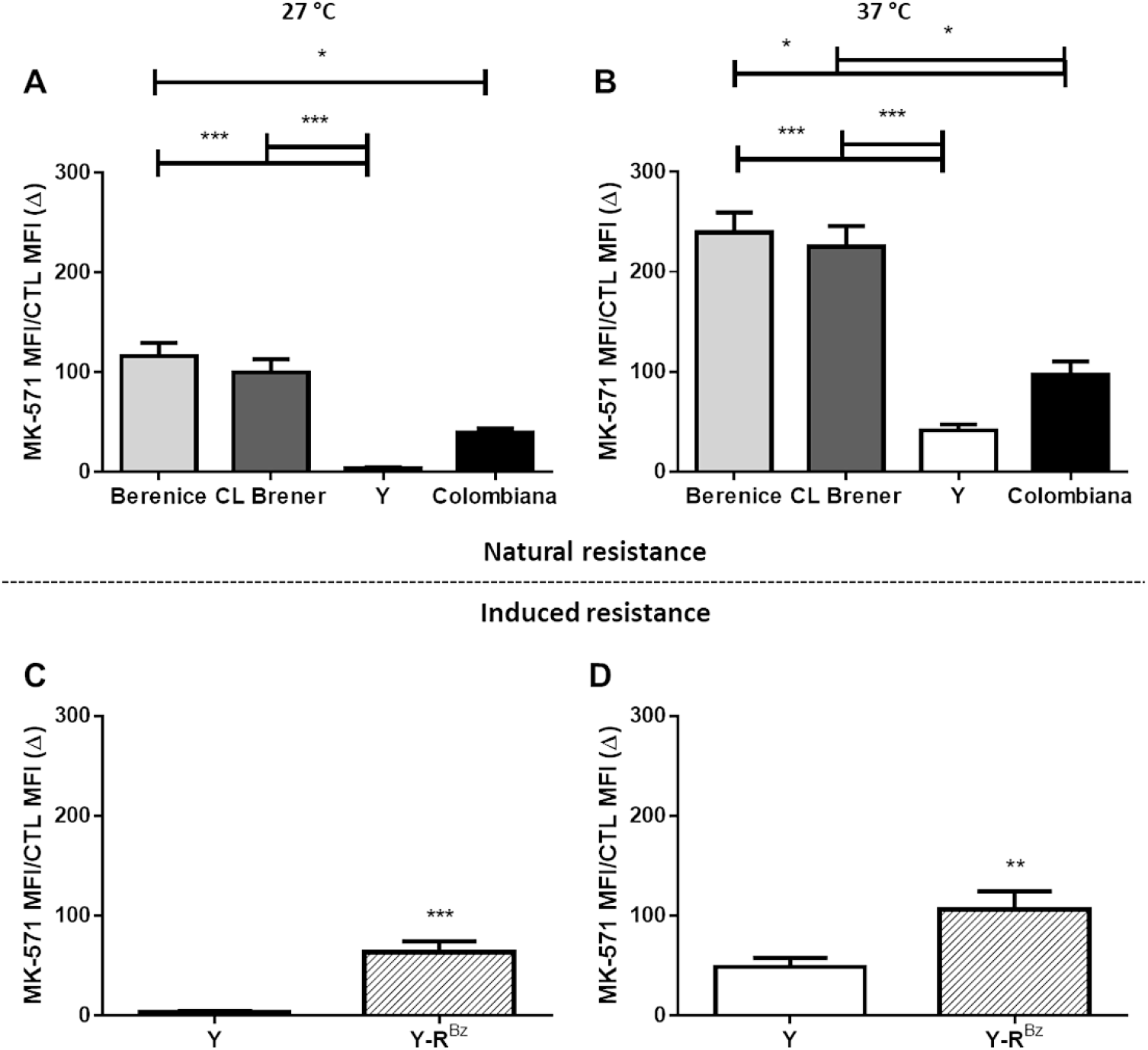
Inhibition indexes (Δ) of CF efflux for MK-571 in *T. cruzi* strains. Δ values represent the ratio of the median fluorescence intensity (MFI) of carboxyfluorescein (CF) of parasites in presence of 200 μM MK-571 to the one on its absence (CTL). Graphs summarize Δ values for efflux assays performed at 27 °C (A and C) and at 37 °C (B and D) in epimastigote forms from CL Brener, Berenice, Colombiana and Y strains and Y-R^BZ^parasites. Bars represent the mean + SEM and the values of significance were represented by (*) for p <0.05, (**) p <0.01 and (***) p <0.001, n=9-10 independent experiments.

### Increased ABCC-like mediated efflux after acquired resistance to benznidazole

Prolonged chemotherapy protocols often lead cells to a drug-adapted phenotype (13), culminating in therapeutic failure and disease relapsing as observed in parasitic infections. Although ABCC-mediated efflux is not related to natural resistance, exposure to benznidazole would select parasites with higher capacity for ABCC transport if it were necessary for survival in this stress condition. For that reason, we evaluated CF efflux in the Y-R^BZ^ parasites. The protocol employed for *in vitro* selection of resistant parasites led to increases in both CF MFI and percentages of CF-positive parasites in presence of MK-571 (Fig. 6G and 6H) and indomethacin (Fig S8G and S8H). Likewise, increasing the assay temperature positively influenced the CF extrusion (Fig. S7G and S7H). Given that nearly all parasites were inhibited by MK-571, results indicate that Y-R^BZ^ population performs ABCC-like mediated efflux. Furthermore, Δ values were higher for the same inhibitor in the selected parasites than in the parental strain (Fig. 7C and 7D), suggesting that ABCC could participate in the acquired resistance to the drug. Remarkably, Y-R^BZ^ were more resistant to the reduction in thiol levels by benznidazole and hemin when compared to parental parasites (Fig. 2B).

### Resistance to benznidazole is not directly dependent on ABCC-like mediated efflux

Aiming to assess whether benznidazole could be directly transported by ABCC-like proteins, the drug was utilized as competitive inhibitor for CF efflux in epimastigote forms from Y strain. Our results suggest otherwise, considering that high concentrations of benznidazole were not able to increase CF MFI nor percentages of CF-positive parasites (Fig. S9). Nevertheless, previous results suggest the formation of thiol-conjugated metabolites after preincubation with benznidazole. As such, the effect of either ABCC inhibition or thiol depletion on the viability of parasites treated with the drug was investigated. Epimastigote forms from Y strain were evaluated by vital staining with propidium iodide (PI). Results indicate that cotreatment with MK-571 and benznidazole was unable to sensitize the parental strain (Fig. 8A); conversely, this protocol reduced the percentage of viable Y-R^BZ^ parasites only at the highest concentration of benznidazole (Fig. 8B). The Y-R^BZ^ parasites were less sensitive than the parental to the MK-571 inhibitor, in which the viability reduction was about 18% against 6% of the resistant parasites. However, the higher concentration of benznidazole alone resulted in a reduction in the percentage of viable cells by 23.33%, which in the presence of MK-571, dropped to nearly 40%. Furthermore, Y strain was susceptible to subtoxic benznidazole concentrations in the presence of buthionine sulfoximine (BSO), an irreversible GSH biosynthesis inhibitor (Fig. 8C). Y-R^BZ^ parasites were further sensitized when an already toxic benznidazole concentration was employed along with BSO (Fig. 8D). BSO had no effect on the viability of the parasites, however, in combination with benznidazole, promoted a 60% reduction for the parental strain, regardless of the concentration of the drug used. In the Y-R^BZ^ strain, cotreatment of BSO with 3 mM benznidazole reduced viability by about 70%. According to results, ABCC participates in the acquired resistance to benznidazole, possibly by efflux of a thiol-conjugated metabolite from benznidazole. Nevertheless, its efflux does not seem to be the dominant pathway of resistance, in view of the inhibition of GSH biosynthesis seemingly showing greater impact on effects of the drug in either parental or resistant parasites.

**Figure 8.**
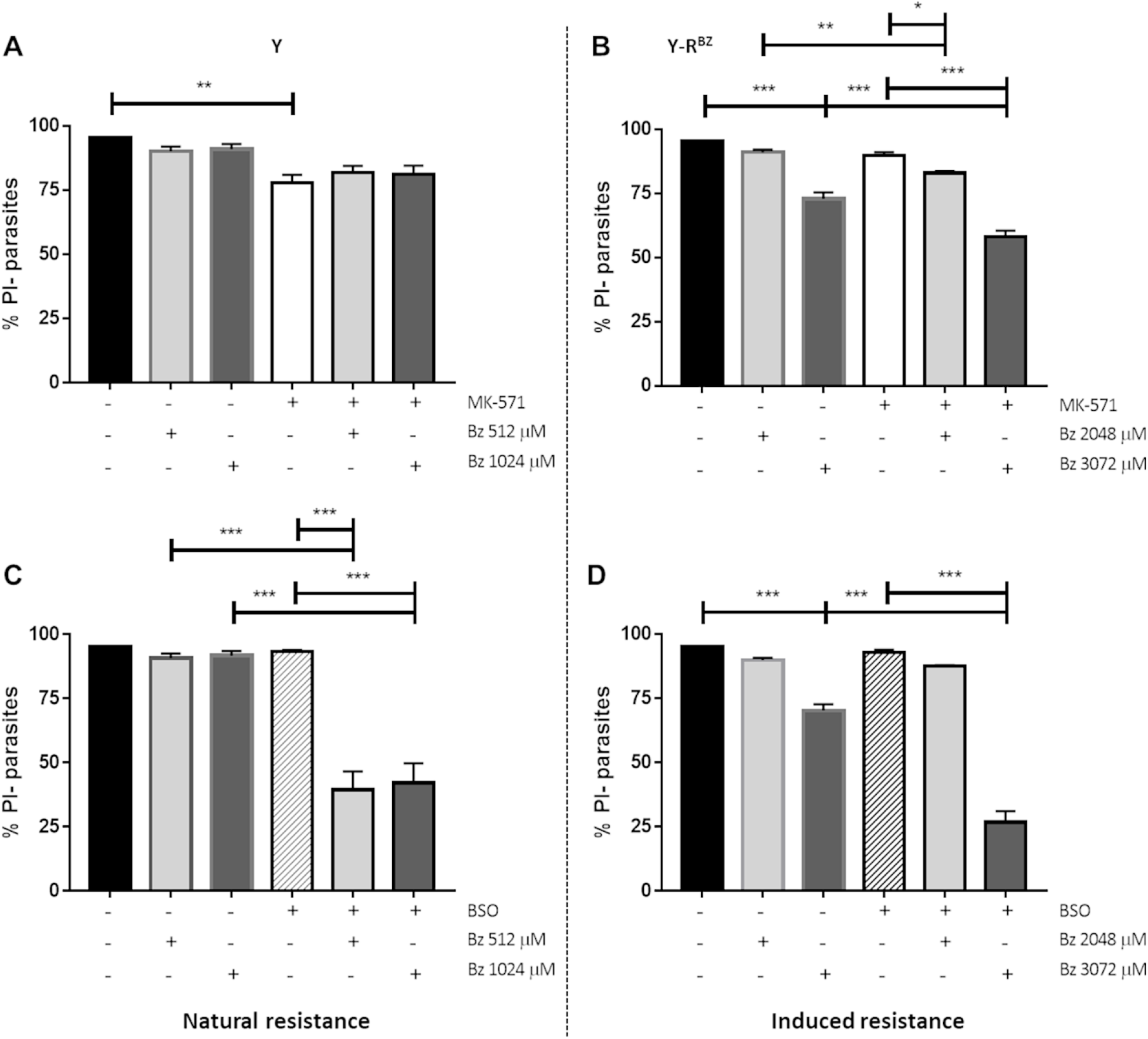
Thiol depletion decreases viability of parasites treated with benznidazole. 10^7^ epimastigotes/mL were incubated in complete medium in absence or presence of benznidazole (ranging from 512 to 3072 μM) alone or in combination with 3 mM buthionine sulfoximine (BSO) or 200 μM MK-571 for 24 h at 27 °C. Parasites were centrifuged and incubated in PBS containing 1 μg/mL propidium iodide (PI) for 15 min. Next, parasites were kept on ice until acquisition by flow cytometry. Percentages of PI-parasites (viable cells) for Y (A and C) and Y-R^BZ^ (B and D). Bars represent the mean + SEM and values of significance were represented by (*) for p<0.05, (**) p<0.01 and (***) p<0.001, n=9 (Y) and n=7 (MK-571/Y-R^BZ^) and n=5 (BSO/ Y-R^BZ^) independent experiments.

## Discussion

Chagas disease poses challenges for chemotherapeutic interventions, as the protozoan has a complex life cycle, diverse mechanisms of evasion from the immune system and high genetic and biochemical diversity (39). Benznidazole is the standard drug for chemotherapy of the disease, although it is inefficent in the chronic phase and in older individuals, and associates to serious side effects making it not indicated to administer it to pregnant patients (40). In addition, the protozoan may show natural resistance to chemotherapy (11) or acquire it due to its prolonged administration (13), evoking the need to research how parasites adapts to stress inducers.

Response pathways to cell stress are essential for the survival of *T. cruzi* in different microenvironments as well as for the emergence of the chemotherapy resistance phenotype. Most cells are able to adapt to stress-inducing stimuli, such as cytokines (e.g. TNF-α, IL-1β and Fas ligands), injury-inducing agents (as bacteria and viruses), cytotoxic agents (heavy metal, chemotherapy and pro-oxidants) and environmental stressors (ultraviolet and ionizing radiation, heat shock and removal of growth factors). Many of these pathways involve the participation of antioxidants such as GSH, the prime defense pathway towards reactive species in general (41), lipid mediators such as ceramides, regulating a number of biochemical and cellular responses (42) and transporters such as ABC proteins, involved in the transport of molecules (43).

Transporters from ABCC subfamily are mainly responsible for the efflux of molecules conjugated to or in cotransport with thiol, as observed in protozoans as *Leishmania* and *Plasmodium* (27) and in mammals (24). In our previous work, we identified an ABCC-like transport in trypomastigote and epimastigote forms of *T. cruzi* Y strain (31). Although we demonstrated that the efflux of fluorescent molecules conjugated to thiol were inhibited by indomethacin, this inhibitor is not specific, being reported in the literature as ABCB1 inhibitor as well (44). In the current work, we demonstrated that both GSH and GSSG acted as competitive inhibitors of ABCC-like mediated transport, promoting the accumulation of CF fluorescent substrate. Remarkably, the reduced form promoted greater modulation of transporter than oxidized as observed by the Δ values, suggesting higher affinity.

Epimastigote forms of *T. cruzi* are exposed to a large amount of heme due to the volume of blood ingested by female triatomines. Heme is a toxic molecule and promotes the formation of ROS, leading to the oxidation of lipids, proteins and nucleic acids (16). In addition, as a lipophilic anion, it inserts itself in phospholipid membranes, leading to leakage as a result of changes in its permeability and selectivity (45). Taken together, these evidences represent a selective pressure that should have been counteracted by protection adaptations of the parasite. Considering that the epimastigote forms of Y strain showed high ABCC-like activity, we evaluated whether ABCC could participate in this protection against iron-induced toxicity. Similar to results observed in *Leishmania* species (46), ABCC-like could be a metal-thiol transporter, since hemin competitively inhibited the CF transport. However, inhibition was slightly higher after depleting of free thiol levels, possibly owing to the production of ROS by the Fenton reaction. In the experiments, we use hemin (Fe^+3^ protoporphyrin IX) as a stress-inducing agent, which can be reduced to heme (Fe^+2^ protoporphyrin IX) in reactions involving superoxide, GSH or ascorbate (47).

ROS are formed as byproducts of stress stimuli but also from normal metabolic processes and affect the metabolism of sphingolipids including ceramides, sphingomyelin and sphingosine (21). In mammals, ceramides act as a coordinator of stress responses, considering that many stress inducers promote accumulation this sphingolipid, usually from sphingomyelin hydrolysis or *de novo* biosynthesis pathway, but often as a result of inhibition of the clearance of ceramides (48). In *T. cruzi*, the *de novo* biosynthesis pathway is similar to that of mammals up to ceramide, which is employed for the synthesis of inositolphosphoceramides (IPC), present in free glycosylinositolphospholipids (GIPL) or in glycoproteins anchoring (49). The synthesis and degradation of IPC are catalyzed by IPC synthase and inositolphospholipid phospholipase C (ISC), which present higher similarity with sphingomyelin synthase and sphingomyelinase of mammal cells respectively (50). IPC can also be hydrolyzed by phosphoinositide phospholipase C found in the surface of parasite (51). In spite of mammal cells, there are not many studies relating sphingolipids as signaling molecules in protozoan parasites, much less in response to cellular stress. Particular works suggest that the presence of ceramides in the plasma membrane has important roles for regulation of processes such as differentiation from trypomastigote to amastigote forms (52), and escape from immune surveillance of the host (53), considering that their anchoring could associate to active shedding into the medium as the case with Ssp4 and *trans*-sialidase glycoproteins (50). Free GIPL are the main glycoconjugates found on the surface of the parasite and related to pathogenicity. The ceramide portion was able to induce apoptosis in activated macrophages (54) and to suppress T cell activation in mammal cells (55). Additionally, GIPL from *T. cruzi* and *T. rangeli* decreased the production of nitric oxide in the insect’s salivary glands, being this an important initial step for parasite survival (56).

Despite the absence of data regarding sphingolipids as mediators of intracellular signaling in *T. cruzi*, it is possible that stress induced by cytotoxic agents such as benznidazole or heme could promote accumulation of ceramides in the parasite in a similar way to mammal cells. Early accumulation could occur via remodeling of IPC, possibly induced by GSH depletion that inhibits sphingomyelinases (57). These enzymes are not described in *T. cruzi*, however they show higher similarity with ISC, which has sphingomyelinase activity as shown for *S. cerevisae* ISC (58). This hypothesis can be supported by other protozoan parasites such as *P. falciparum*, in which stress induced by the chemotherapeutic agents artemisinin and mefloquine induced ceramide accumulation in a GSH-dependent manner (59). Another alternative route would be derived from *de novo* biosynthesis, seeing that drugs as Tamoxifen inhibited IPC synthase in *L. amazonensis* (60).

ABC proteins transport lipids, including phospho- and sphingolipids (61). ABCC1 was directly implicated to the transport of sphingosine-1-phosphate (25), glucosylceramide and sphingomyelin in mammalian strains (26) in a GSH-dependent manner. Assuming that ceramides could be induced in response to stress by cytotoxic agents in *T. cruzi*, we evaluated the role of ABCC-mediated transport in the accumulation of this sphingolipid. Preincubation with sphingosine, employed to stimulate *de novo* biosynthesis of endogenous ceramides, promoted CF accumulation in epimastigote forms of Y strain. In order to evaluate the possibility of direct transport of ceramides, C6-NBD-cer were employed in the efflux assay. Indeed, we observed that the transport of C6-NBD-cer was inhibited by MK-571 as well as by the depletion of ATP and free thiol levels, demonstrating for the first time that ceramides are direct substrates of ABCC.

ABCC-like transporters could have evolved as adaptive advantages for prevention of death signaling and as promoters of asymmetric arrangement in the plasma membrane. It is unknown how sphingolipids carry on intracellular signaling, considering that they are found in the outer leaflet of the lipid bilayer (50). Nonetheless, spontaneous exchanges between the two leaflets of the lipid bilayer can occur quickly for phospholipids without high polar headgroups such as ceramides (62). In the present study, we used synthetic short-chain ceramides that are membrane permeable allowing for its accumulation in the cytosol. Regardless, transport of long chain lipids could occur as well, considering that ABC proteins are required for the transport of cuticular wax (C ≥ 26) in plants (63) and that ABCB1 reconstituted in membranes was able to transport fluorescent long-chain phospholipids (64). It is worth noting that the transport mediated by ABC proteins may differ depending on the nature of the substrate. Amphipathic molecules would be taken from the cytosolic side and extruded into the extracellular medium, while hydrophobic molecules would be flipped to the outer leaflet or translocated to another membrane (65).

Knowing that ABC proteins are overexpressed in the MDR phenotype, we decided to evaluate whether the ABCC-like activity could relate to cell protection in naturally or in benznidazole-induced resistant strains. To estimate natural resistance, three *T. cruzi* strains were evaluated for ABCC-like transport and compared to Y strain analyzed in our previous work (31). CL Brener and Berenice were employed as models of naturally sensitive strains while the Y and Colombiana as strains naturally resistant to benznidazole. ABCC activity was inhibited by MK-571 in all strains, however, the naturally sensitive strains showed higher Δ values, indicating higher activity and possibly, ABCC expression. For this reason, it does not seem that ABCC is involved with the natural resistance of *T. cruzi* strains, and other intrinsic factors should explain the Y and Colombiana resistant phenotype. On the other hand, Y strain was used in order to mimic the resistance acquired by prolonged exposure to benznidazole, since it showed the lowest Δ value among all strains. After treatment, Y-R^BZ^ parasites showed a high IC_50_ value for the drug and an increase in ABCC-mediated efflux when compared to the parental. ABCB1-like activity does not influence natural or acquired resistance, in view of that no strain presented Rho 123 efflux, suggestive of absence of protein expression, at least in plasma membrane. As such, an ABCC-like transporter could mediate benznidazole efflux. Results suggest the opposite, because the drug was not able to competitively inhibit the CF transport even in high concentrations.

Benznidazole is a prodrug and its activation produces reactive species capable of associating with free thiols. Consequently, Y-R^BZ^ parasites showed resistance to reducing thiol levels either by drug or hemin administration, suggesting an adaptation of the GSH biosynthesis pathway in the phenotype of acquired resistance. According to this evidence, the parasites’ viability was analyzed after treatment with combinations of benznidazole and an ABCC transport inhibitor (MK-571) or an irreversible inhibitor of the GSH biosynthesis (BSO). Inhibition of transport had no impact on the effect of benznidazole on the Y or Y-R^BZ^ parasites, except in the latter when a toxic concentration was employed. The reduction of the toxic effect of MK-571 on the Y-R^BZ^ parasites compared to the Y strain corroborates the increased activity on selected parasites. Treatment with benznidazole and BSO demonstrated that GSH has a relevant impact for the response to drug-induced stress. Y strain showed a marked reduction in viability at non-toxic concentrations of the drug with BSO. Y-R^BZ^ parasites tolerate the stress of GSH depletion and benznidazole to a certain extent; subsequently the combined treatment reduced the viability to levels comparable to the parental, reversing the resistant phenotype. The present data suggest that ABCC is necessary for the extrusion of toxic benznidazole metabolites, most likely in conjugation to or cotransport with GSH. In addition, benznidazole is reduced by a type I nitro-reductase in *T. cruzi*, resulting in the generation of the glyoxal cytotoxic metabolite that can interact with free thiol (38).

Faundez *at al*. demonstrated that the thiol biosynthesis was important for resistance to chemotherapy, once treatment with BSO had increased the toxicity of benznidazole and nifurtimox, another chemotherapeutic possibility for Chagas’ disease (66, 67). Studies have shown that antioxidant enzymes such as tryparedoxin peroxidase (68) and iron-superoxide dismutase-A (69) participate of protection to cope with reactive species from benznidazole. In contrast, participation of transporters involved in the response to xenobiotics has been minimally explored. Other ABC transporters could be involved in the resistance phenotype, considering that *T. cruzi* presents 27 ABC genes. ABCG1 was found to be overexpressed in strains naturally sensitive to benznidazole (70, 71), but its functionality has not been studied so far. In *Leishmania*, ABCG1 and ABCG2 are transporters of thiol and phosphatidylserine, implicated in oxidative metabolism, autophagy, metacyclogenesis and infectivity (72), observations that highlight possible roles for this transporter for *T. cruzi* that should be evaluated in future studies. In conclusion, we believe that our results bring light to processes of *T. cruzi* adaptation to natural or synthetic stresses, in which efflux transporters such as ABC proteins and markedly the subfamily ABCC, act performing the role they accomplish best: reducing cytotoxicity by efflux of xenobiotics and managing levels of mediators of cell death in coordination with the GSH pathway.

## Experimental procedures

### Cultures of Trypanosoma cruzi strains

CL Brener, Berenice and Colombiana strains of *T. cruzi* were gently donated by Dr. Policarpo A. Sales Junior of the Rene Rachou Research Center of the Oswaldo Cruz Foundation from Minas Gerais, Brazil. The Y strain was kindly donated by Prof. Celio Freire-de-Lima from the Institute of Biophysics Carlos Chagas Filho (IBCCF) of the Federal University of Rio de Janeiro (UFRJ), Rio de Janeiro, Brazil.

Epimastigote forms were cultivated at 27 °C in Brain and Heart Infusion medium (BHI, BD Biosciences, São Paulo, SP, Brazil) supplemented with 10% fetal bovine serum (FBS, Life Technologies of Brazil, São Paulo, SP, Brazil), 20 μg/mL folic acid (Sigma-Aldrich, São Paulo, SP, Brazil), 12.5 μg/mL hemin (Sigma-Aldrich) and 50 μg/mL gentamicin (Sigma-Aldrich). For subcultures, epimastigote forms were collected weekly and 10^6^ parasites/mL were suspended in complete BHI medium. All centrifugations were performed at 1,000×g for 10 min at room temperature.

### MTT reduction assay

The tetrazolium salt 3-(4,5-dimethylthiazol-2-yl)-2,5-diphenyltetrazolium bromide, also known as MTT, was used to assess mitochondrial reducing activity. During respiration, cells convert the water-soluble MTT to the insoluble purple product formazan, which is solubilized in DMSO and its concentration determined by optical density. Briefly, 10^6^ epimastigotes/mL were distributed on 96-well culture plates in complete BHI medium, and benznidazole was added to final concentrations of 1 to 1024 μM. After 48 h, plates were centrifuged and the supernatants discarded, followed by addition of PBS supplemented with 2 g/L glucose and 10% FBS. Plates were incubated for 4 h at 27 °C in the dark after adding a MTT/PMS solution (2.5 mg/mL of MTT and 0.22 mg/mL PMS (both from Sigma-Aldrich). Next, plates were centrifuged and the supernatants discarded. Formazan crystals were dissolved in DMSO (Sigma-Aldrich) and plates were shaken for 20 min at room temperature protected from light. Absorbance reading was performed at 570 nm on a Beckman Coulter AD340 spectrophotometer (Beckman Coulter, Brea, CA, USA) (73). Experiments were performed in triplicate and IC_50_ of mitochondrial reducing activity after exposure to benznidazole were determined by logarithmic regression of the normalized percentage curve using the GraphPad Prism software (version 7.0, GraphPad, San Diego, CA, USA). Benznidazole was provided by Dr. Nubia Boechat Andrade of Institute of Technology in Pharmaceuticals of the Oswaldo Cruz Foundation from Rio de Janeiro, Brazil.

### In vitro induction of resistance to benznidazole

10^6^ epimastigotes/mL of the Y strain were maintained in complete BHI medium with the addition of 30 μM benznidazole (IC_50_ obtained after MTT assays) at 27 °C. After 2 days, the parasites were centrifuged, supernatants discarded and complete BHI medium added to allow replication of the surviving parasites for the next 5 days. Parasites were submitted to the same procedure up to 8 weeks. Then, the concentration of benznidazole was increased in 10 μM per week up to a final concentration of 120 μM (74). Resistance to benznidazole was then evaluated by the MTT reduction assay as described before. Thereafter, the benznidazole-adapted Y strain was named Y-R^BZ^. For subcultures, 10^6^ epimastigotes/mL were suspended in complete BHI medium containing 120 μM benznidazole for 7 days.

### ATP depletion

The irreversible inhibition of glyceraldehyde-3-phosphate-dehydrogenase by alkylating agent as IAA is able to reduce glycolysis and, consequently, ATP levels (75). Then, 10^7^ epimastigotes/mL were incubated in absence or presence of 2 mM IAA or (Sigma-Aldrich) in PBS for 1 h at 27 °C. Next, parasites were centrifuged, supernatants discarded and parasites suspended in PBS for the ABC-mediated efflux assay. In this case, PBS was employed instead of RPMI during the efflux assay due to absence of glucose.

### ABC-mediated efflux assay

The efflux assay was divided into 30 min steps of accumulation and efflux of substrate (76). For the ABCC-mediated efflux assay, the CFDA dye (Life Technologies of Brazil) was employed as precursor. In the cytosol, CFDA is hydrolyzed and originates the fluorescent substrate CF (77), which is transported to extracellular medium by ABCC subfamily members (78). Briefly, 10^7^ epimastigotes/mL were incubated with 50 μM CFDA diluted in RPMI medium (Sigma-Aldrich) in absence (control) or presence of 600 μM indomethacin or 200 μM MK-571 as ABCC inhibitors (both from Sigma-Aldrich). After the accumulation step, parasites were centrifuged and then suspended in RPMI medium in absence or presence of the inhibitors. After the efflux step, parasites were centrifuged, supernatants discarded and parasites suspended in PBS supplemented with 5% FBS and kept on ice for immediate acquisition by flow cytometry. Alternatively, 5 mM GSH, 5 mM GSSG, 100 or 200 μM hemin (all from Sigma-Aldrich) or 0.5, 1, 2 or 3 mM benznidazole were employed as competitive inhibitors in the same way as ABCC inhibitors. Otherwise, parasites were preincubated with hemin or sphingosine (Sigma-Aldrich) for 1 h and removed before the assay. In this case, no inhibitor or modulator was added to efflux assay.

For evaluation of ceramide efflux by ABCC transporters, 10 μM C6-NBD-cer (Avanti Polar Lipids, Alabaster, AL, USA) was employed as fluorescent substrate in absence or presence of 200 μM MK-571 in RPMI.

Similarly, the naturally fluorescent substrate Rho 123 was employed to analyze ABCB1-mediated efflux assay. Therefore, epimastigote forms were incubated with 100 nM Rho 123 diluted in RPMI medium in absence or presence of 50 μM CsA, 10 μM VP (Sigma-Aldrich) or 10 μM TFP (Sigma-Aldrich) as ABCB1 inhibitors in the accumulation and efflux steps. CsA was kindly donated by Dr. Marcia Capella from the IBCCF, UFRJ, Rio de Janeiro, Brazil.

The efflux assays were performed at 27 °C (insect vector temperature) and/or 37 °C (temperature of positive control). As negative control, parasites were not exposed to dyes. MDR cells Lucena-1 or FEPS were employed as positive control due to overexpression of ABCB1 and ABCC1 respectively (79–81). These cells were gently offered by Dr. Vivian Rumjanek from the Institute of Medical Biochemistry Leopoldo de Meis, UFRJ, Rio de Janeiro, Brazil.

### Determination of intracellular thiols

10^7^ epimastigotes/mL were incubated in the absence or presence of 512 or 1024 μM benznidazole or 200 μM hemin in PBS supplemented with 2 g/L glucose for 3 h at 27 °C. After, parasites were centrifuged, supernatants discarded and parasites incubated in PBS containing 1.5 μM CMFDA at 27 °C for 15 min prior to acquisition by flow cytometry (82). CMFDA is an acetoxymethyl ester derivative, being able to cross the plasma membrane efficiently. In the cytosol, it reacts with exposed sulfhydryl radicals, forming the compound TMF. As negative control, parasites were not exposed to dye. As positive control of thiol depletion, parasites were incubated in PBS containing 100 μM NEM, an alkylating agent, for 1 h and removed before adding of CMFDA.

### Assessment of cellular viability

Staining of nonviable parasites was performed by the DNA intercalation dye PI, which is readily excluded by live cells with intact membranes (83). 10^7^ epimastigotes/mL were incubated at 27 °C for 24 h with concentrations ranging from 1 to 3072 μM benznidazole diluted in complete BHI medium in absence or presence of 200 μM MK-571 or 3 mM BSO, an irreversible inhibitor of GSH biosynthesis. Parasites were then centrifuged, supernatants discarded and parasites suspended in 1 μg/mL PI diluted in PBS and incubated for 15 min prior to acquisition by flow cytometry. As positive control of cell death, cells were incubated with distilled water for 30 min before addition of PI. As autofluorescence control, parasites were not exposed to dye.

### Flow cytometry analyses

The CF, Rho 123, TMF and C6-NBD-cer fluorescence intensities were acquired on the FL1-H channel (530/30 bandpass filter) while PI fluorescence on the FL3-H channel (670LP filter) of a FACSCalibur (BD Biosciences, San Jose, CA, USA). Post-analysis was performed in the software Summit (version 4.3, Dako Colorado, Fort Collins, CO, USA) on at least 10,000 viable cells, gated in accordance with forward (FSC) and side scatter (SSC) parameters representative of cell size and granularity. MFI data and percentages of parasites were acquired from histograms for each dye. A negative/low fluorescence gate was designed containing 95% of control parasites from the histogram origin while a high fluorescence gate contained the remaining. The Δ value in the CF efflux assay was calculated by the ratio of MFI of inhibited cells to MFI of control cells.

### Statistical analysis

Statistical analyzes were performed using the software GraphPad Prism. For two comparisons nonpaired, the t-student or Mann-Whitey tests were respectively employed for parametric and nonparametric data. For more than two comparisons, one-way ANOVA or Kruskal-Wallis tests were respectively employed for parametric and nonparametric data. For paired nonparametric data, the Wilcoxon or Friedman tests were performed for two or more than two comparisons. The Turkey’s, Sidak’s, Dunn’s or Dunnet’s post-tests were used according to the compared columns. Significance values were represented by (*) for p<0.05, (**) for p <0.01 and (***) for p <0.001.

## Data availability

All the data are contained within the article and Supporting Information.

## Acknowledgements

We thank Dr. Celio Freire-de-Lima and Dr. Policarpo A. Sales Junior for offering *T. cruzi* strains. We are grateful Dr. Nubia Boechat Andrade, Dr. Vivian Rumjanek and Dr. Marcia Capella for donating benznidazole, MDR cells and CsA, respectively.

## Funding and additional information

This work was supported by grants from Conselho Nacional de Desenvolvimento Científico e Tecnológico - CNPq, Fundação Carlos Chagas Filho de Amparo à Pesquisa do Estado do Rio de Janeiro - FAPERJ and Coordenação de Aperfeiçoamento de Pessoal de Nível Superior - CAPES.

## Footnotes

The abbreviations used are: Δ: inhibition index of transport ABC: ATP-binding cassete ABCB1: subfamily B, member 1/ also known as P-glycoprotein ABCC: subfamily C/ also known as MRP ABCG1: subfamily G, member 1 ABCG2: subfamily G, member 2/ also known as BCRP BCRP: Breast Cancer Resistance Protein BHI: brain heart infusion BSO: buthionine sulfoximine BZ: benznidazole C6-NBD-ceramide: N-[6-[(7-nitro-2-1,3-benzoxadiazol-4-yl)amino]hexanoyl]-D-erythro-sphingosine) CF: carboxyfluorescein CFDA: 5(6)-carboxyfluorescein diacetate CMFDA: 5-chloromethylfluorescein CsA: cyclosporin A FBS: fetal bovine serum GIPL: glycosylinositolphospholipids GSH: reduced glutathione GSSG: glutathione disulfide form/ oxidized glutathione IAA: iodoacetic acid IPC: inositolphosphoceramide ISC: inositolphospholipid phospholipase C MDR: multidrug resistance MK-571: (E)-3-[[[3-[2-(7-chloro-2-quinolinyl)ethenyl]phenyl][[3-(dimethylamino)-3-oxopropyl]thio]methyl]thio]-propanoic acid MFI: median fluorescence intensity MRP: multidrug resistance protein/ABCC MTT: 3-(4,5-dimethylthiazol-2-yl)-2,5-diphenyltetrazolium bromide NEM: N-ethylmaleimide PGP: P-glycoprotein/ ABCB1 PI: propidium iodide PMS: phenazine methosulfate Rho 123: rhodamine 123 ROS: reactive oxygen species T(SH)_2_: trypanothione TFP: trifluoperazine TMF: thiol-conjugated methylfluorescein VP: verapamil

